# *Plasmodiophora brassicae* infection threshold – how many resting spores are required for infection of canola (*Brassica napus*)

**DOI:** 10.1101/2021.05.21.445124

**Authors:** Kher Zahr, Yalong Yang, Alian Sarkes, Snezana Dijanovic, Heting Fu, Michael W. Harding, David Feindel, Jie Feng

## Abstract

Clubroot, caused by *Plasmodiophora brassicae*, is an important disease of canola and other brassica crops. Improved understanding of host and pathogen biology is frequently useful in guiding management strategies. In order to better understand infection thresholds, seven-day old seedlings of canola cultivar Westar were inoculated with serially diluted concentrations of *P. brassicae* resting spores. Controlled soil and plant inoculation assays were performed and the plants maintained in a greenhouse for 42 days and clubroot disease severity evaluated visually. Clubroot symptoms were observed in soils containing as low as one spore/mL soil and on plants inoculated with as few as ≤ 100 resting spores. These thresholds were lower than any previously reported. The results indicated the importance of highly sensitive detection methods for *P. brassicae* diagnosis and quantification methods for clubroot risk prediction in soils. Furthermore, these results highlighted the low probability of obtaining *P. brassicae* single spore isolates.

## Introduction

Clubroot, caused by the protist *Plasmodiophora brassicae* Woronin, is an important threat to Canadian canola (*Brassica napus*) production (Hwang et al. 2012). In the Canadian Prairies, clubroot was first identified on canola in 2003, in a dozen fields near Edmonton, Alberta (Tewari et al. 2005). The disease has since then spread throughout central Alberta (Strelkov et al. 2015) and has also been confirmed in canola fields in Saskatchewan (Ziesman et al. 2019), Manitoba (Froese et al. 2019), Ontario (Al-Daoud et al. 2018) and North Dakota (Chittem et al. 2014). *Plasmodiophora brassicae* can produce large numbers of resting spores, which can survive in the soil for up to 20 years (Wallenhammar 1996). Resting spores are the primary source of inoculum in natural conditions, thus, direct and indirect measurements of resting spores in soil have been extensively used for *P. brassicae* detection and diagnosis, and forecasting of clubroot risk.

Many methods have been developed for clubroot diagnosis, among which PCR-based assays such as end-point PCR, quantitative PCR (qPCR) and digital PCR are the most sensitive and accurate (Faggian and Strelkov 2009). The sensitivities of the PCR diagnosis from soil were generally reported at 1000 spores/g soil for PCR (Cao et al. 2007), 500-1000 spores/g soil for qPCR (Wallenhammar et al. 2012; Li et al. 2013) and 100 spores/g soil for digital PCR (Wen et al. 2020).

In 2016 and 2019, soil samples containing serial dilutions of resting spores were provided to plant health/diagnostic labs across Canada to evaluate their efficiency for PCR-based diagnosis of clubroot (J. Feng, unpublished). In both years, the efficiencies from the labs were similar to those previously reported so that all labs reported *P. brassicae* detection from 1000 spores/g soil samples, but this concentration was also the lower limit of detection for most labs. It was believed that the lower limit of detection was comparable with the lowest concentration of resting spores in soil that could cause visible clubroot (Donald and Porter 2009). At the concentration of 1 × 10^3^ spores/g soil, *P. brassicae* may cause infection, but no aboveground symptoms and little impact on crop yield could be observed (Kageyama and Asano 2009).

These results indicated that many spores (i.e. 1 × 10^3^ spores/g soil) on or in the vicinity of a root hair would be required to establish an infection and subsequently generate clubroot galls. In other words, 1 × 10^3^ spores/g soil appeared to be near the lower limit concentration for clubroot establishment. With this infection threshold in mind, trying to obtain a single spore isolate of *P. brassicae* would be extremely difficult, or perhaps impossible. Previous studies reported that the efficiency of obtaining single spore isolates was low, as indicated by the ratio between plants with galls and total plants inoculated with single spores, for examples, three out of 450 (0.7%; Jones et al. 1982), eight out of 600 (1.3%; Scott 1985) and two out of 164 (1.2%; Voorrips 1996). Recently, Akarian et al. (2021) reported much higher efficiencies, with 2-22% depending on the origins of the isolates and a summarized efficiency of 8.5% (34 out of 400). However, all the above-mentioned single spore isolates were generated on Chinese cabbage. No single spore isolate has been obtained from canola.

With the illogicality between the perceptions that the establishment of clubroot infection in field conditions needs many spores, and that a single spore can cause infection and subsequently generate clubroot galls in the creation of single-spore isolates, we conducted this greenhouse study to investigate the clubroot infection thresholds with serious dilutions of *P. brassicae* resting spores in soil or on plant roots. The objectives were to 1) find the lowest spore concentration in soil that could cause visible clubroot symptoms on canola, 2) measure the possibility/efficiency of infections caused by single spores and 3) determine the sensitivity of *P. brassicae* detection required to accurately estimate disease risk in soil.

## Materials and methods

### Host plants and *Plasmodiophora brassicae* populations

Clubroot-susceptible canola cultivar Westar was used as the host plant exclusively in this study. Two *P. brassicae* populations, one pathotype 3H and the other composed mainly of pathotype 5x, were mixed at the ratio 1:1 and the mixture was used to prepare inocula. Origins of these two populations, as well as population maintenance and gall mixture preparation followed Zahr et al (2021). Resting spore suspensions were prepared according to Zhang et al. (2015) from samples of the gall mixture.

### Soil inoculation assay

The spore suspension was adjusted to 1 × 10^8^/mL, from which further dilutions were prepared. Two-L square plastic pots were filled at 1.5 L per pot with soilless mix (Sunshine mix #4, Sun Gro Horticulture, Vancouver, BC, Canada). The pots were soaked in water for 5 hours and then kept on greenhouse benches overnight to drain the excess water. The weight of 1.5 L soilless mix at this time point was 1464.4 g ± 33.3 (mean ± SD, n = 10). The soilless mix from ten pots was poured into a 46-L concrete mixer. With the mixer rotating at 35 rpm, the soilless mix was inoculated with resting spores by spraying 150 mL of a spore suspension using a plastic spray bottle. After inoculation, the mixer kept rotating at 35 rpm for 10 min. The inoculated soilless mix was used to re-fill the ten pots. The inoculation was conducted with a set of 10× serial dilutions of resting spore from 1 × 10^8^ spores/mL to 1 × 10^2^ spores/mL, which resulted in ten pots of inoculated soilless mix at each of the seven concentrations from 1 × 10^6^ spores/mL soilless mix to 1 spore/mL soilless mix. In addition, ten pots of soilless mix inoculated with 150 mL water were also prepared. The inoculations were conducted with the order from low spore concentrations to high spore concentrations. Immediately after inoculation, ten seedlings of Westar, generated from seeds on moist filter paper at 24°C and 16-h photoperiod for seven days, were transplanted into each pot. The pots were placed in individual plates on greenhouse benches as randomized complete blocks (n = 10) with one block occupying one bench and the eight pots in each block being 2 feet away to each other. The greenhouse was maintained at 24°C/18°C (day/night) with a 16-h photoperiod. Starting from the third day after inoculation (dai), the pots were watered from the bottom every second day with tap water at pH 6.4 (adjusted with HCl). This assay was conducted twice.

### Plant inoculation assay

Small pieces of filter paper (7 mm in diameter) were prepared using a paper punch. One piece of the filter paper was placed into the cap of a 1.5-mL microcentrifuge tube that had been cut off from the tube. The filter paper was misted with 25 μL water and a canola seed was placed at the centre of the paper. The caps were incubated in a transparent plastic bag at 24°C with a 16-h photoperiod. After seven days, a set of 10× serial dilutions of resting spore suspensions was prepared, from 1 × 10^7^ spores/mL to 1 × 10^3^ spores/mL. The root of the seedling in each cap was inoculated with 1 μL of spore suspension taken from one of the serial dilutions (Fig. 1). One μL water was used as the negative control treatment. After inoculation, the caps were kept in the transparent plastic bag at 24°C with a 16-h photoperiod. After 48 hours, the seedlings were transplanted into 500-mL plastic pot filled with soilless mix (Sunshine mix #4) with one plant per pot. The transplanting was done by picking up the filter paper on which the seedling was situated and burying the lower part of the stem, the root and the filter paper into the soilless mix. After transplanting, the pots were placed 1 foot apart on individual plates on greenhouse benches in a completely randomized design. Greenhouse conditions and watering were identical as the soil inoculation assay. This assay was conducted three times. In each experiment, for all treatments (spore concentrations) except 1 × 10^3^ spores/mL, 30 plants were inoculated. For the treatment of 1 × 10^3^ spores/mL, 180, 170 and 165 plants were inoculated in the first, second and third experiments, respectively.

**Fig 1.**
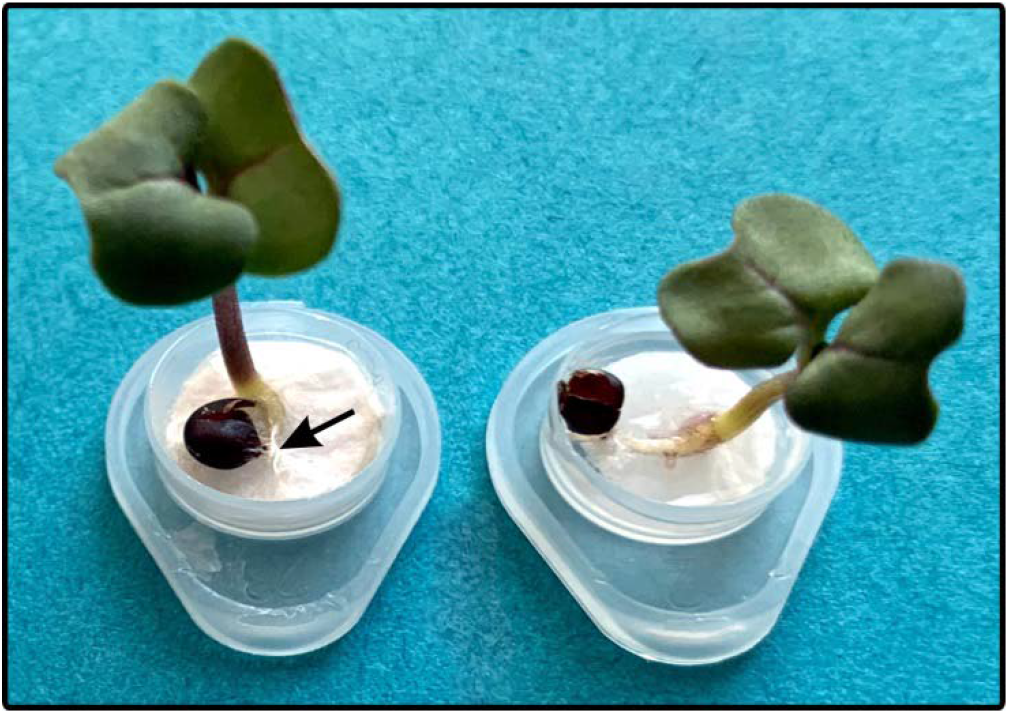
Inoculation of seven-day-old canola seedling with 1 μL *Plasmodiophora brassicae* resting spores. The arrow indicates the inoculated root.

### Spore counting

In each of the three plant inoculation experiments, a 40-mL aliquot was taken from the 1 × 10^3^ spores/mL inoculum right before inculcation and kept at 4°C for less than 3 hours. In a 50-mL centrifuge tube, the aliquot was centrifuged at 4000 rcf for 10 min. After remove the supernatant, the pellet was dissolved in 4 mL water, making a spore suspension at 1 × 10^4^ spores/mL. Using a hemocytometer (Fisher Scientific Canada, Ottawa, ON; cat number: 02-671-6) bearing two 0.1-μL counting chambers, spore number in 0.1-μL samples was counted 300 times for the first experiment and 100 times for the second and the third experiments under a Leica DM500 microscope (Leica Microsystems Canada, Concord, ON), with 12 μL sample applied on the chamber for each counting.

### Clubroot rating

For the soil inoculation assay, clubroot severity on each plant was evaluated at 42 dai using a 0- to-3 scale (Kuginuki et al. 1999), where 0 = no clubbing, 1 < one-third of the root with symptoms of clubbing, 2 = one-thirds to two-thirds clubbed, and 3 > two-thirds clubbed. Severity ratings on all the ten plants in each pot were converted to an index of disease (ID) using the formula of Strelkov et al. (2006). For the plant inoculation assay, the presence or absence of clubroot galls on each plant were visually investigated at 42 day after transplanting.

### Statistics

All statistics was done using Microsoft Excel. In the soil inoculation assay, data from each experiment was subjected to analysis of variance (ANOVA). Based on the ANOVA results, differences between treatments (spore concentrations in the soil) were assessed with Tukey’s multiple comparison test (P ≤ 0.05) using the Excel add-in DSAASTAT developed by Dr. Onofri at the University of Perugia, Italy (http://www.casaonofri.it/repository/DSAASTAT.xls). In the spore counting assay, calculation of Poisson distribution and Chi square test were conducted with the POISON.DIST function and the CHISQ.TEST function, respectively.

## Results

### Soil inoculation assay

In the two repeated experiments of the soil inoculation assay, tiny galls, rated as severity scale 1, were observed on plants in the treatment of 1 spore/mL soil (Fig. 2A and B), with two plants in experiment 1 (in alternative pots) and one plant in experiment 2. In both experiments, large galls were observed on plants in the treatments of ≥10 spores/mL soil (Fig. 2C). No gall was observed in the treatment of water control (Fig. 2D). The disease index (ID) curves from the two experiments were similar (Fig. 3). In both experiments, there were significant differences between the ID of < 1000 spores/mL soil treatments and those of ≥ 1000 spores/mL soil treatments. These results indicated that clubroot infection can occur when the spore concentration is at 1 spore/mL soil, albeit at a low frequency, and that 1000 spores/mL soil is a key spore concentration threshold for appearance of visible clubroot disease symptoms.

**Fig. 2.**
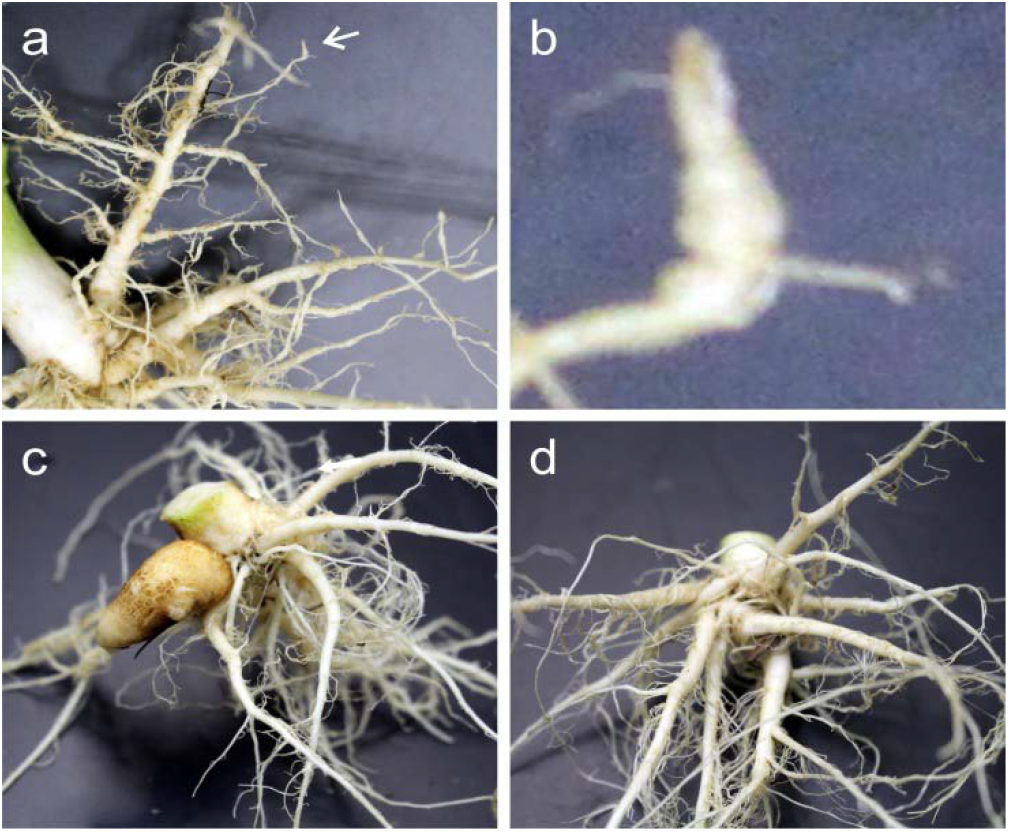
Galls developed on canola 42 days after growing in soil inoculated with *Plasmodiophora brassicae* resting spores at the final concentration of 1 spore/mL soil (a and b) and 10 spores/mL soil (c) or inoculated with water as the negative control (d). A tiny gall in (a) is indicated by an arrow and this part of (a) is magnified in (b).

**Fig 3.**
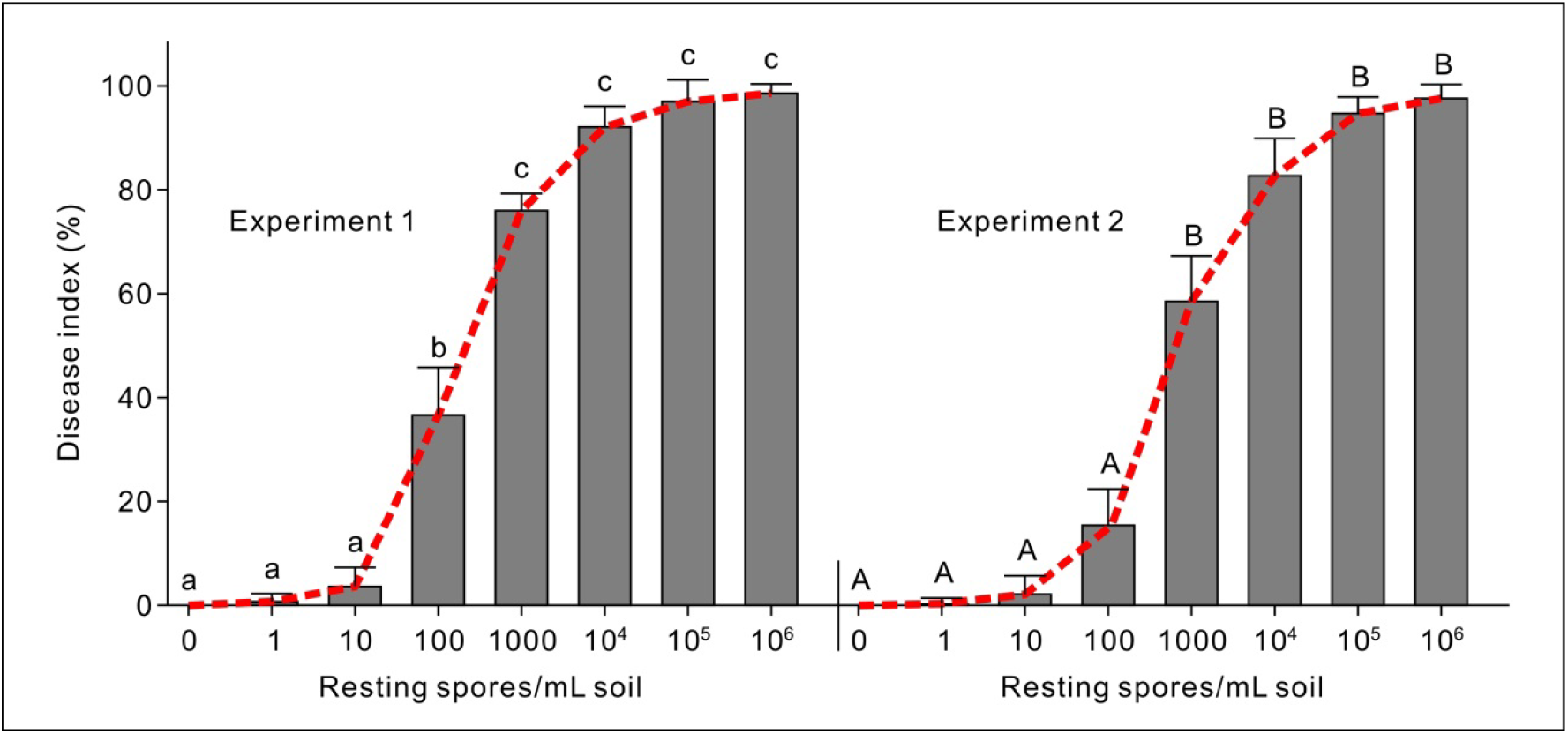
Disease index of clubroot developed on canola 42 days after growing in soil that was inoculated with *Plasmodiophora brassicae* resting spores at final concentrations from 0 to 1 × 10^6^ spores/mL soil. Means in the plots topped by the same letter do not differ based on Tukey's multiple comparison test at P ≤ 0.05 (n = 10).

### Single spore counting

At very low concentrations, the distribution of particles in aliquots follows a Poisson distribution (Ahrberg et al. 2018). We calculated the possibilities of obtaining certain numbers of spores in a 1-μL aliquot from a 1 spore/μL solution (Table 1). The possibility of obtaining ≥ 1 spore would be 63.21% (1-36.79%). To verify that the number of spores in the 1-μL samples taken from the 1 × 10^3^ spores/mL inoculum followed the Poisson distribution, we counted the spores 300 times for the first and 100 times for the second and the third inoculation experiment. Since the hemocytometer was designed for counting spore numbers in a volume of 0.1 μL, all countings were conducted after concentrating the 1 × 10^3^ spores/mL inoculum to 1 × 10^4^ spores/mL. The three counting experiments showed similar patterns to those of the calculated Poisson distribution, all with Chi square values supporting goodness of fit (Table 1). This result indicated that in our plant inoculation assay, approximately 63% of the plants treated with 1 μL of the 1 × 10^3^ spores/mL suspension were inoculated by one or more spore(s).

**Table 1.** Counting of *Plasmodiophora brassicae* resting spores in 0.1-μL aliquots taken from resting spore suspensions at a concentration of 1 × 10^4^ spores/mL

| Spores per count | Expected ratio (%)* | Experiment 1 |  | Experiment 2 |  | Experiment 3 |  |
| --- | --- | --- | --- | --- | --- | --- | --- |
|  |  | Observed | Expected | Observed | Expected | Observed | Expected |
| 0 | 36.79 | 114 | 110 | 35 | 37 | 31 | 37 |
| 1 | 36.79 | 112 | 110 | 33 | 37 | 38 | 37 |
| 2 | 18.39 | 55 | 55 | 23 | 18 | 20 | 18 |
| ≥3 | 8.03 | 19 | 24 | 9 | 8 | 11 | 8 |
| Chi square |  | 1.220 |  | 1.747 |  | 2.189 |  |
| Two-tailed *P* value |  | 0.7482 |  | 0.6264 |  | 0.5341 |  |

\*Ratios of obtaining the corresponding spore numbers if the countings conform to Poisson distribution.

### Plant inoculation assay

In the three plant inoculation experiments, 515 plants were inoculated with a 1-μL aliquot of spore suspension at the concentration of 1 × 10^3^ spores/mL (Table 2). A single gall was observed from one of the 515 plants (Fig. 4). A single gall was also observed from one of the 90 plants and one of the 89 plants inoculated with 1-μL aliquots of a spore suspension at the concentration of 1 × 10^4^ spores/mL and 1 × 10^5^ spores/mL, respectively. In contrast, galls, either single or multiple, were observed on 11 and 33 plants inoculated with 1-μL aliquots of a spore suspension at the concentration of 1 × 10^6^ spores/mL and 1 × 10^7^ spores/mL, respectively. These results indicated that the infection efficiency was very low when the spore number was less than 1000.

**Table 2.** Inoculation of canola roots with 1-μl aliquots of serially diluted concentrations of *Plasmodiophora brassicae* resting spores

| Spores | Experiment 1 |  | Experiment 2 |  | Experiment 3 |  | In Total |  |
| --- | --- | --- | --- | --- | --- | --- | --- | --- |
|  | Plant | Gall | Plant | Gall | Plant | Gall | Plant | Gall |
| 0 | 29 | 0 | 30 | 0 | 30 | 0 | 89 | 0 |
| 1 | 170 | 0 | 180 | 1 | 165 | 0 | 515 | 1 |
| 10 | 30 | 0 | 30 | 1 | 30 | 0 | 90 | 1 |
| 100 | 30 | 0 | 30 | 0 | 29 | 1 | 89 | 1 |
| 1000 | 29 | 3 | 30 | 4 | 30 | 4 | 89 | 11 |
| 10000 | 30 | 10 | 30 | 12 | 30 | 11 | 90 | 33 |

**Fig. 4.**
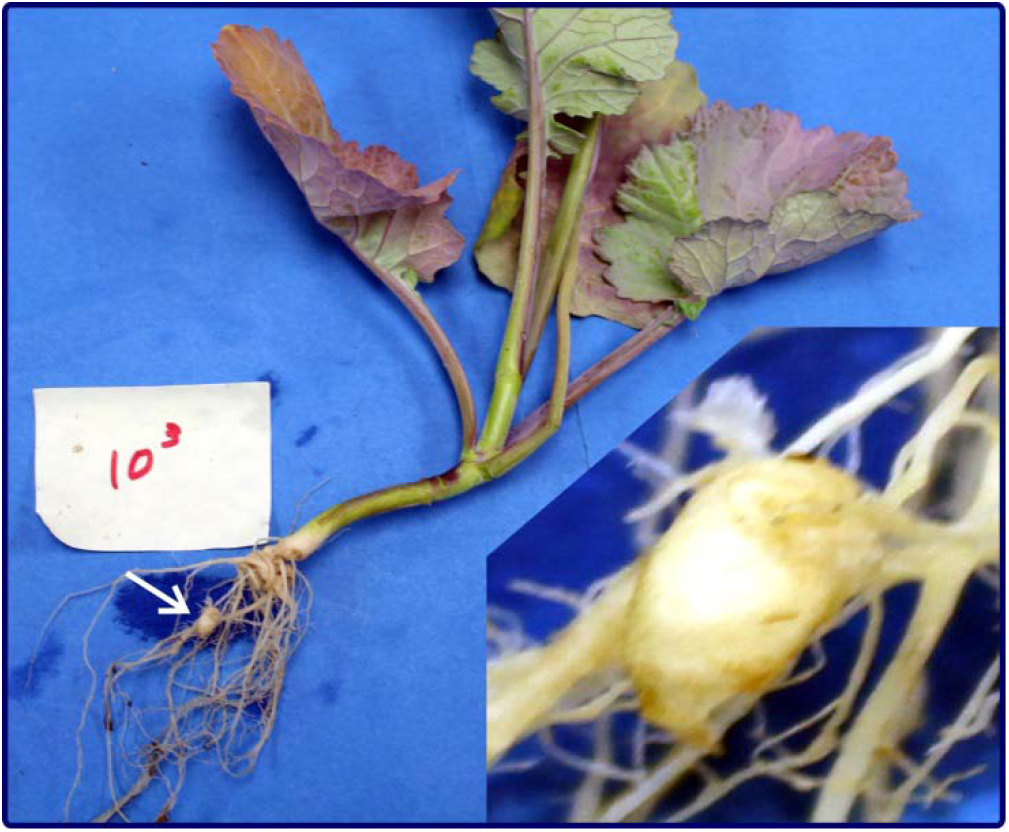
A gall developed on a canola plant inoculated with 1 μL *Plasmodiophora brassicae* resting spores at a concentration of 1 × 10^3^ spores/mL and transplanted into soil 48 hours after inoculation. Photo was taken at 42 day after transplanting. The area indicated by the arrow is enlarged in the lower right.

## Discussion

The results from the soil inoculation assay indicated that 1 spore/mL (approximately 1 spore/g) of soil was sufficient to obtain visible clubroot symptoms. Currently, the lower limit of PCR-based *P. brassicae* detection from soil is approximately 1000 spores/g soil. This finding revealed a potentially urgent need of more sensitive *P. brassicae* detection methods for soil samples. Detection thresholds lower than 1000 spores/g soil have occasionally reported, for example, by using digital PCR. However, we would hesitate to use a PCR or qPCR system that reported detection of *P. brassicae* from a spore concentration as low as 1 spore/g soil. This is because if DNA was extracted from a soil sample containing 1 spore and then only a small portion (e.g. 1 out of 50 μL) of the resultant DNA was used as the template in a PCR reaction, it would be almost impossible to amply a single-copy DNA fragment. Furthermore, if a DNA sample extracted from 1000 resting spores was used to make serial dilutions and subsequent PCR or qPCR amplifications could produce positive signal from the 1000× dilution, it should not be concluded that the protocol could detect the pathogen in a soil or root sample containing one spore. This is because the efficiency of DNA extraction is affected by both the plant or soil background and the proportion of target in the sample mixture. The efficiency of a PCR detection system is not only dependent on the PCR reaction per se, but also significantly influenced by sample preparation method and how the DNA was extracted. For example, one might extract DNA from 100 mg of soil and another might first purify the resting spores from 50 g of the soil sample and then extract DNA from the purified spores.

For a PCR-based detection method, regardless of the reaction sensitivity, at least one copy of the target DNA fragment must be present in the template. Most commercial DNA extraction kits recommend using less than 200 mg starting material for DNA extraction and dissolving the resultant DNA in 50-100 μL water or the provided buffer, which would dilute the copy number of the target fragment present in the template. Thus, we proposed the following points one should consider when developing and advocating new PCR-based methods for *P. brassicae* detection: 1) try to target a DNA region with multiple copies in the genome, 2) define what sample was used for DNA extraction, i.e., from purified spores or directly from soil and 3) indicate the lowest number of spores from which the DNA in one positive reaction was derived. These points should also be considered when interpreting results from previously published PCR-based detection or quantification studies.

In this study we had initially tried to create single spore isolates for *P. brassicae*, but it was extremely difficult to confirm the presence of a single spore in an inoculum sample. Not only was it practically impossible to see a single spore on any solid surface under a microscope, but also with spores suspended in water, the lack of contract, and the movement/vibration of the spores, made the single-spore determination very challenging. These circumstances led to ambiguous results and unfruitful, yet time consuming, evaluations. Furthermore, when only one or a few particles are present in a droplet, it is difficult to differentiate resting spores from starch grains or other debris of similar size. Due to these complications, we bypassed the step of single spore isolation and utilized the calculated probabilities of obtaining single spores based on Poison distribution. Our results from the spore counting assay indicated that the 1-μL aliquots taken from the 1 × 10^3^ spores/mL suspension could be used to assess the efficiency of single spore isolation.

Although we obtained one gall from the 515 plants inoculated by 1 μL of the 1 × 10^3^ spores/mL suspension, only one gall was obtained from 90 and 89 plants inoculated by 1 μL of 1 × 10^4^ spores/mL and 1 × 10^5^ spores/mL suspensions, respectively. There are two possible reasons for this low efficiency: 1) a *P. brassicae* infection may need more than one spore or 2) single-spore infection on canola is rarer compared to that on Chinese cabbage. The low infection efficiencies of the inocula containing less than 1000 spores, along with the difficulties visualizing single spores within a spore suspension, had prevented the generation of single-spore populations of *P. brassicae*, not only in this study, but also in others (J. Feng unpublished data). It also became evident that there is currently no way to demonstrate that an obtained population was single-spore derived because without verification of single spore inoculum, resulting root galls may result from infection by two or more spores that are genetically identical.

We could not directly compare our results from the soil inoculation assay and the plant inoculation assay because there was no way to measure how many spores in the soil contributed to the observed infection. The microenvironment in soil will influence resting spore germination and zoospore infection in a very different way than those directly applied to the root surface.

Nevertheless, the results of both assays demonstrated that the inoculum concentrations in soil and on plant required for the establishment of visible clubroot symptoms is lower and higher, respectively, than previously thought.

In summary, we concluded from this study that 1) in 1 spore/mL soil *P. brassicae* is able to cause clubroot, and as a consequence the sensitivity of *P. brassicae* detection and quantification needs to be increased and 2) the efficiency of *P. brassicae* single spore canola root infections is very low.

## Funding information

Financial support was received from Canadian Agricultural Partnership (no. 601322).

## Compliance with ethical standards

## Conflict of interest

The authors declare that they have no conflict of interest.

## Research involving human and/or animals

No animals or data from human participants were involved in this study.

## Notes

### Competing Interest Statement

The authors have declared no competing interest.

